# Revealing a hidden intermediate of rotatory catalysis with X-ray crystallography and Molecular simulations

**DOI:** 10.1101/2021.12.07.471682

**Authors:** Mrinal Shekhar, Chitrak Gupta, Kano Suzuki, Takeshi Murata, Abhishek Singharoy

**Affiliations:** School of Molecular Sciences, Arizona State University, Tempe, AZ, USA; Center for Development of Therapeutics, Broad Institute of MIT and Harvard, Cambridge, MA, USA; Department of Chemistry, Graduate School of Science, Chiba University, Inage-ku, Chiba, Japan; Membrane Protein Research and Molecular Chirality Research Centers, Chiba University, Inage-ku, Chiba, Japan

## Abstract

The mechanism of rotatory catalysis in ATP-hydrolyzing molecular motors remain an unresolved puzzle in biological energy transfer. Notwithstanding the wealth of available biochemical and structural information inferred from years of experiments, knowledge on how the coupling between the chemical and mechanical steps within motors enforces directional rotatory movements remains fragmentary. Even more contentious is to pinpoint the rate-limiting step of a multi-step rotation process. Here, using Vacuolar or V_1_-type hexameric ATPase as an exemplary rotational motor, we present a model of the complete 4-step conformational cycle involved in rotatory catalysis. First, using X-ray crystallography a new intermediate or ‘dwell’ is identified, which enables the release of an inorganic phosphate (or P_i_) after ATP hydrolysis. Using molecular dynamics simulations, this new dwell is placed in a sequence with three other crystal structures to derive a putative cyclic rotation path. Free-energy simulations are employed to estimate the rate of the hexameric protein transformations, and delineate allosteric effects that allow new reactant ATP entry only after hydrolysis product exit. An analysis of transfer entropy brings to light how the sidechain-level interactions transcend into larger-scale reorganizations, highlighting the role of the ubiquitous arginine-finger residues in coupling chemical and mechanical information. Inspection of all known rates encompassing the 4-step rotation mechanism implicates overcoming of the ADP interactions with V_1_-ATPase to be the rate-limiting step of motor action.

## INTRODUCTION

V-type ATPase from *Enterococcus hirae* is a prototypical ATP-driven rotary molecular motor. It harnesses energy from ATP hydrolysis to pump ions across biological membranes^1^. Crystallographic studies reveal that V-ATPases possess an overall three-dimensional structure, composed of a hydrophilic domain (V_1_) and a membrane-embedded, ion-transporting domain (V_o_) connected by central and peripheral stalk(s)^2–8^. The V_1_ domain consists of an A_3_B_3_ ring (composed of 3 repeats of A and B subunits) and a central stalk **(Fig. 1)**. Ubiquitous to all rotary ATPases, including the famous F_1_-ATP synthase, V_1_-ATPases partake in ‘rotatory catalysis’_1_– a catalytic mechanism, wherein the chemical reactions e.g. ATP hydrolysis occurs through conformational rotation of the A_3_B_3_ ring and physical rotation of the central stalk^2,3^.

**Fig. 1:**
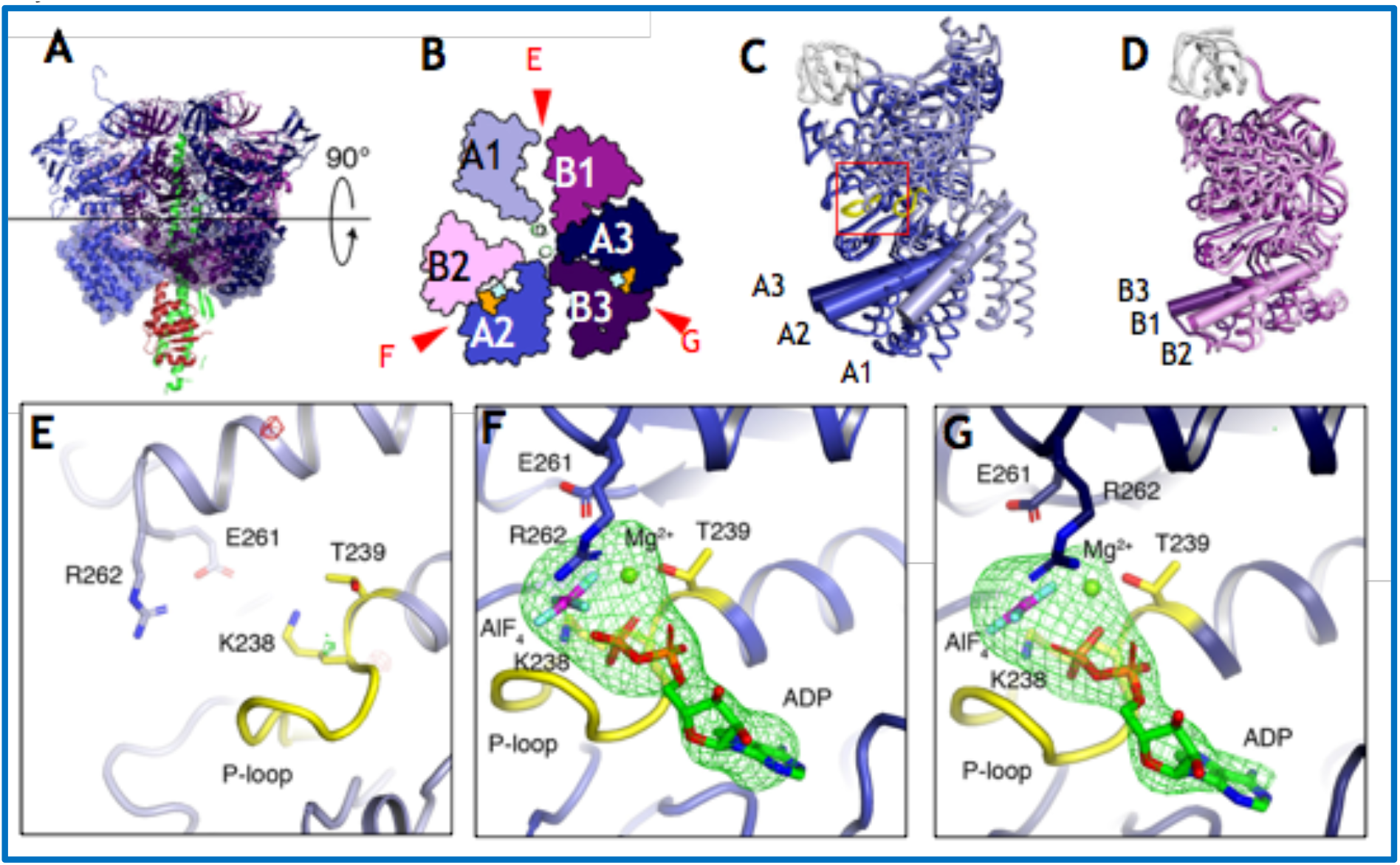
Structure of 2_(ADP · AlF4)_V_1_-bound V_1_ complex. **(A)** Side view of 2_(ADP · AlF4)_V_1_. **(B)** Top view of the C-terminal domain (shown in A at transparent surface) of 2_(ADP · AlF4)_V_1_from the cytoplasmic side. Red arrows indicate the nucleotide-binding sites. The bound ADP and AlF_4_^-^ molecules are shown in space-filling representation and colored orange and cyan, respectively. **(C-D)** Superimposed structure at the N-terminal β-barrel region (white) of three structures of A subunits (C) and B subunits (D) in 2_(ADP · AlF4)_V_1_. A subunits are colored light blue (A1 or A^e^), dark blue (A2 or A^b^), and darker blue (A3 or A^t^) in order of openness, and similarly for the B subunits: dark purple (B1 or B^e^), light purple (B2 or B^b^) and darker purple (B3 or B^t^). The P-loops are shown in yellow. **(E-G)** Magnified view of the nucleotide-binding sites of in 2_(ADP · AlF4)_V_1_, corresponding to the red box of panel C. The positions of the nucleotide-binding sites correspond to the symbol written on panel B. The |Fo|-|Fc| maps calculated without ADP:Mg^2+^ and aluminum fluoride molecules at the binding pockets contoured at 4.0 sigma are shown in red (negative) and green (positive).

The rotatory catalysis mechanism was originally proposed by Boyer in the 1980s^9^, and Walker and colleagues have spent the following two decades to find key structural intermediates (popularly known as *dwell* states) along the ATP hydrolysis pathway^10–15^, which is also supported by Senior’s mutational and biochemical analysis of the motor^16^. However, molecular details of a contiguous reaction pathway linking the individual dwell states are being uncovered only recently by using a combination of fast single-molecule experiments^17^, and multiscale molecular dynamics simulations^18–20^. Contributing to this body of information, we have determined the crystal structures of the so-called “catalytic dwell”, “ATP-binding dwell”, and “ADP-release dwell” of the motor, and proposed the V_1_-ATPase rotational mechanism model based on the crystal structures and molecular simulations^21,22^.

Illustrated in Fig. S1, our proposed multi-site catalytic mechanism includes: (*i*) ATP hydrolysis into ADP and inorganic phosphate (or P_i_) in the *tight* AB domains of the A_3_B_3_ ring within the catalytic dwell [PDB: 3VR6]^2^, accompanied with a loosening of the *tight* AB interface and straightening of the DF stalk subunit; (*ii*) after the hydrolysis reaction, the ADP remains bound to the *tight* interface, and the neighboring *empty* AB domains opens up to become *bindable* for accepting a new ATP in the pocket, transforming to the ATP-binding dwell [PDB: 5KNB]^3^; (*iii*) the new ATP binds to this bindable site making it *half-closed* and the motor transforms to the ADP-release dwell [PDB: 5KNC]^3^; (*iv*) following ADP release, the DF stalk undergoes a deformation and subsequent 120° rotation, so the symmetry of the system is reset back to the catalytic dwell [PDB: 3VR6], completing one cycle of rotatory catalysis^3^.

The hydrolysis-product release step is implicated as rate-determining in other related motors, such as the hexameric helicases^23^ and even in F-type ATPases^18^. But the crystallographic structures of V_1_-ATPase determined so far did not include states immediately after ATP hydrolysis and before P_i_ release, keeping details of this key step of the catalytic cycle elusive. In this study, we report the crystal structure of *E. hirae* V_1_ ATPase (EhV_1_), in which the aluminum fluoride (or AlF_4-_) and ADP molecules are bound at two nucleotide-binding sites. It has been discussed that the aluminum fluoride and ADP bound structure mimics the transition state of ATP hydrolysis. Analogous to the structures of myosin_24_, F_1_-ATPase_12_ and other ATPases that have been obtained with aluminum fluoride bound, we seek the post-hydrolysis mechanism of the V_1_ motor action. Compared with the crystal structure in the catalytic dwell state that has an ATP-bound tight AB pair, we find the model with an ADP · AlF_4_^-^-bound AB pair to be marginally open. This AB interface is open enough to allow P_i_ release but not as wide to allow release of the ADP. We therefore interpret that the ADP · AlF_4_^-^-bound ATPase structure corresponds to a state of waiting for P_i_ release following ATP hydrolysis, and label this state as the “P_i_-release dwell”. Starting from the crystal structure of the catalytic dwell, we performed molecular dynamics and free energy simulations to model the P_i_ release pathway following ATP hydrolysis. An allosteric mechanism is described that connects product release from the tight pocket with increased ATP affinity in the neighboring bindable pocket. This way, using simulations we place the P_i_-release dwell in the sequence of events lining the overall rotatory catalysis mechanism in EhV_1_, and complete the first molecular description of the entire conformational cycle joining four X-ray crystallographically determined intermediates. A kinetic analysis of the product release mechanism offers insights on the rate-determining step of molecular motor action.

## RESULTS

In what follows, first, the EhV_1_ with ADP and aluminum fluoride is crystallized, and the 3D-structure of the ADP · AlF_4_^-^ bound V_1_ is determined employing X-ray crystallography. Second, local and global structural differences between this ADP · AlF_4_^-^-bound V_1_ model and that from the ATP-binding catalytic dwell state are computed to ascertain the location of the P_i_-release dwell along the rotatory catalysis cycle **(Fig. 2)**. Third, molecular dynamics simulations reveal a mechanism of P_i_ release, which couples the P_i_-release dwell with the following ATP-binding and ADP-release dwells. Finally, noting that the path ensuing from this ATP-binding dwell to ADP-release dwell resetting back to another catalytic dwell is already established in our previous studies^22,25,26^, a complete model for rotatory catalysis in V-type ATPases is accomplished.

**Fig. 2:**
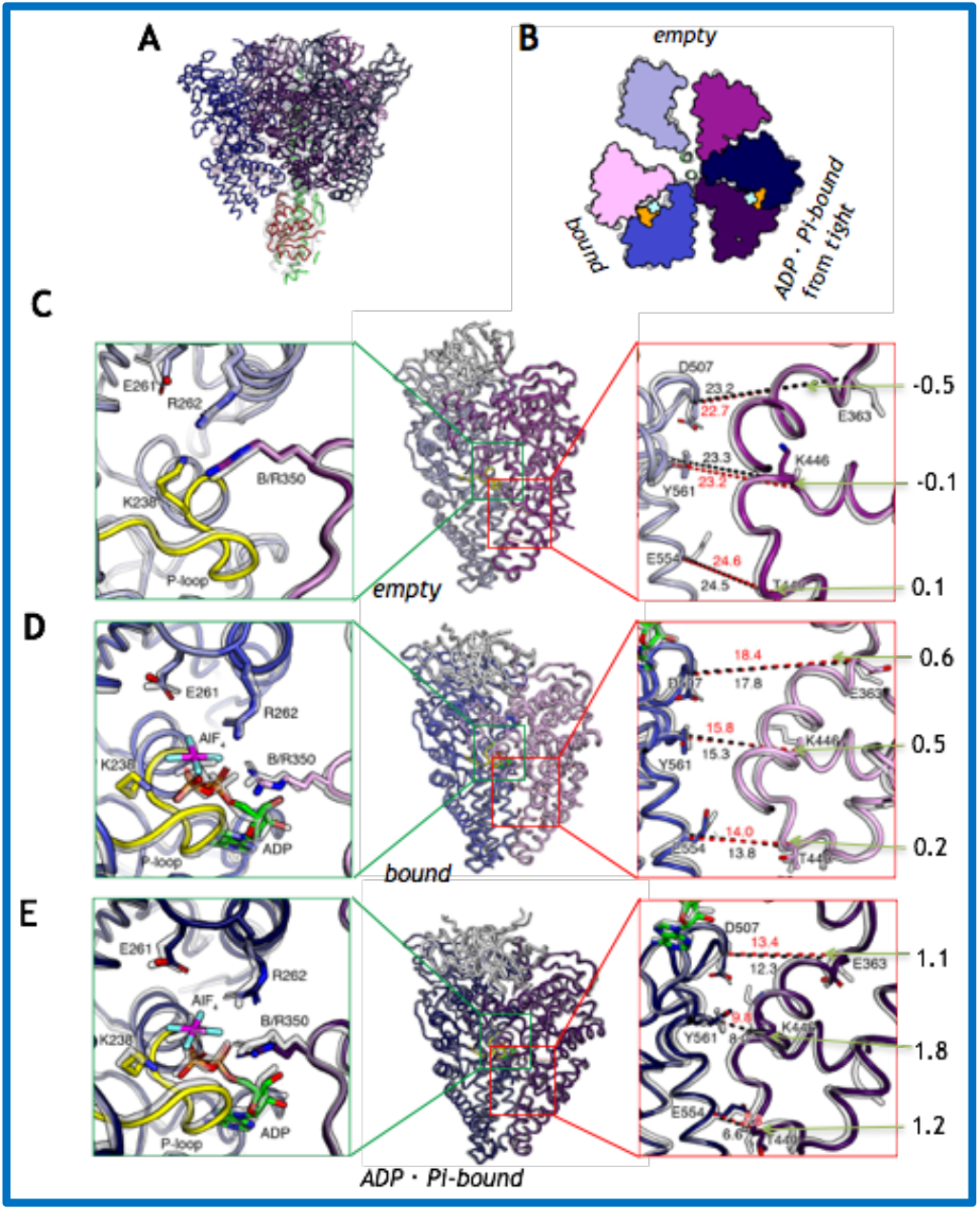
Comparison of the structures of 2ADP · AlF_4-_-bound and 2AMP · PNP-bound V_1_ complexes. **(A-B)** The structures of 2_(ADP · AlF4)_V_1_ (colored) are superposed on the AMP · PNP-‘*bound*’ or 2_(AMP · PNP)_V_1_ conformations (shown in gray). (A) Side view, and (B) top view of the C-terminal domain from the cytoplasmic side. The bound AlF_4_^-^ and ADP molecules are shown in space-filling representation and colored cyan and orange, respectively. **(C-E)** The ‘*empty*’ (C), ‘*bound*’ (D), and ‘*ADP · Pi-bound*’ (E) form in 2_(ADP · AlF4)_V_1_ (colored) are superimposed on those of 2_(AMP · PNP)_V_1_ (shown in transparent gray) at A subunits (residues 67-593). Left panels, magnified views of the nucleotide binding sites, corresponding to the green box of middle panels. The middle panels, side views of AB pairs. Right panels, magnified views of the interface of C-terminal domains, corresponding to the red box of middle panels. Red (2_(ADP · AlF4)_V_1_) and black (2_(AMP · PNP)_V_1_) dotted lines indicate the distances (Å) between C_α_ atoms. The numbers outside the panel represent the value of the lengths of the red dotted lines minus the lengths of the black dotted lines.

### Crystallization of V_1_-ATPase with ADP and aluminum fluoride reveals a new intermediate

It has been reported that the ATPase activity of bovine mitochondrial F_1_-ATPase is inhibited in the presence of ADP and aluminum fluoride, and the crystal structure of ADP · AlF_4_^-^ bound F1 corresponds to a post-hydrolysis (and pre-product release) state in the catalytic cycle^12^. In this study, we purified EhV_1_ in the presence of ADP and AlF_4_^-^, analogous to the case of the bovine F_1_-ATPase (see Methods). The ATP hydrolysis activity of the purified EhV_1_ was not observed, suggesting that the EhV_1_ was inhibited by binding of ADP · AlF_4_^-^ in the nucleotide-binding site(s). We crystallized the inhibited EhV_1_ and obtained a crystal structure at 3.8 Å resolution on an *R* factor of 22.7% and a free *R* factor of 26.6% (**Table 1**).

**Table 1.** Data collection and refinement statistics.

|  |  |
| --- | --- |
| **Data collection** |  |
| Space group | $P2_12_12_1$ |
| Cell dimensions |  |
| $a, b, c$ (Å) | 127.7, 128.9, 231.6 |
| $\alpha, \beta, \gamma$ (°) | 90.0, 90.0, 90.0 |
| Resolution (Å) | 46.13-3.82 (4.05-3.82) |
| $R_{\text{merge}}$ | 0.172 (0.953) |
| $I/\sigma I$ | 7.61 (1.54) |
| Completeness (%) | 99.5 (99.0) |
| Redundancy | 3.9 (3.8) |
| **Refinement** |  |
| $R_{\text{work}}/R_{\text{free}}$ (%) | 22.69/26.56 |
| R.m.s deviations |  |
| Bond lengths (Å) | 0.003 |
| Bond angles (°) | 0.509 |
| Ramachandran plot statistics (%) |  |
| Favored regions | 97.8 |
| Allowed regions | 2.1 |
| Outliers | 0.0 |

### Structure of the V_1_ complex with ADP and aluminum fluoride

The obtained crystal structure was composed of a hexagonally arranged A_3_B_3_ complex and a central axis DF complex as the previously reported structures of EhV_1_^2^ (**Fig.1**A-B). We superimposed the N-terminal β-barrel domain of the three A or B subunits to evaluate the conformational differences in the EhV_1_ complex because the β-barrel domain should be fixed to form an alternatively arranged ring^2^ (**Fig.1**C-D). All A and B subunits showed different conformations, suggesting that this inhibited EhV_1_ is also formed in an asymmetric structure as the other published structures of EhV_1_. Three nucleotide-binding sites are at the interface between the A and B subunits. Two strong electron density peaks were observed in two nucleotide-binding sites (**Fig. 1**E-G). ADP:Mg^2+^ and AlF_4_^-^ molecules were fitted well into both density peaks. Therefore, we denote this structure as 2_(ADP · AlF4)_V_1_ from here on.

Structural differences between 2 ADP · AlF_4_^-^ bound and 2 AMP · PNP bound V_1_ complexes highlight molecular conformations before and after P_i_ release. We previously reported that 2 AMP-PNP-bound EhV_1_ (denoted as 2_(AMP · PNP)_V_1_), corresponding to the catalytic dwell state during rotatory catalysis, consists of three different conformation AB pairs: *empty* (site that cannot bind nucleotides), *bound* (site that can bind ATP), and *tight* (site waiting for ATP hydrolysis). The structures of 2_(ADP · AlF4)_V_1_ and 2_(AMP · PNP)_V_1_ are compared in Fig. 2. The overall structure of 2_(ADP · AlF4)_V_1_ was very similar to that of bV_1_ (root mean square deviation: RMSD = 0.78 Å) (Fig. 2A,B). Especially, the *empty* site of the 2_(AMP · PNP)_V_1_ is very similar to the AB pair with no nucleotide of 2_(ADP · AlF4)_V_1_ (RMSD = 0.51 Å): these nucleotide-binding sites are almost identical, and the distances between the C-terminal domains of the AB pairs are also very similar (**Fig. 2**C). The *bound* site of 2_(AMP-PNP)_V_1_ is also similar to an AB pair which binds ADP:Mg^2+^ and AlF_4_^-^ in 2_(ADP · AlF4)_V_1_ (RMSD = 0.51 Å): these nucleotide-binding sites bind different substrates, but the structures of the binding sites and the distances between the AB pairs are very similar (**Fig. 2**D).

In contrast, the remaining third AB pair bound ADP:Mg^2+^ and AlF_4_^-^ molecules of 2_(ADP · AlF4)_V_1_ shows prominent local differences when compared with the *tight* site bound AMP · PNP:Mg^2+^ of 2_(AMP-PNP)_V_1_, although the overall RMSD between this two AB pairs is 0.62 Å. Illustrated in **Fig. 2**E, we find that the conformations of the conserved residues of E261 and R262 of A subunit and the Arg-finger (R350 in B subunit) are deviated by 1.1–1.8 Å, which is probably induced by binding of AlF_4_^-^ molecule instead of the gamma-phosphate of AMP · PNP; such conformational differences were much lesser (between 0.1-0.6 Å) with the *empty* and *tight* sites. Furthermore, the C-terminal domain of the AB pair shows slightly open conformation that may allow P_i_ release but not as wide to allow the release of the ADP (**Fig. 2**E). From these findings, we designated the AB pair, which was more open than the *tight*, as *ADP · P*_*i*_*-bound* form, and interpreted that the structure corresponds to the state of waiting for P_i_ release (denoted as “P_i_-release dwell”).

### P_i_-release dwell - a new step in the rotational mechanism

Here, we integrate the P_i_-release dwell into the rotational mechanism model for V_1_-ATPase based on all available crystal structures (see Fig. S1 and Fig. 3): The rotation mechanism model starts with a catalytic dwell in which V_1_-ATPase binds two ATP molecules at the bound and tight sites. Since the Arg-finger residue R350 in the *tight* site is in close proximity to the ATP gamma-phosphate, this ATP is waiting for hydrolysis. The cycle begins with the hydrolysis of this ATP (**Fig. 3**A). The ATP in *tight* site is hydrolyzed and produces ADP and P_i_, which induce the conformational change to *ADP · P*_*i*_*-bound* form, waiting for P_i_ release from the binding site; this state represents the P_i_-release dwell in **Fig. 3**B. The product, P_i_, is released from the slightly open conformation of this *ADP* · *P*_*i*_*-bound* site. Then the resulting *ADP-bound* ATPase, complexing with just one ADP is created. Herein, a conformational transition of the *empty* site (120° apart from the ADP-bound site) renders it to a *bindable* form. The *empty* site has a low affinity to bind nucleotides^2^; however, due to this conformational change, a new ATP becomes accessible to the *bindable* form. This structure is, therefore, referred to as the “ATP-binding dwell”, waiting for new ATP binding (**Fig 3**C) to subsequently enable the rotation of the central stalk (Fig. S1)^8^. In the following we evaluate the proposed model with molecular dynamics or MD simulations.

**Fig. 3:**
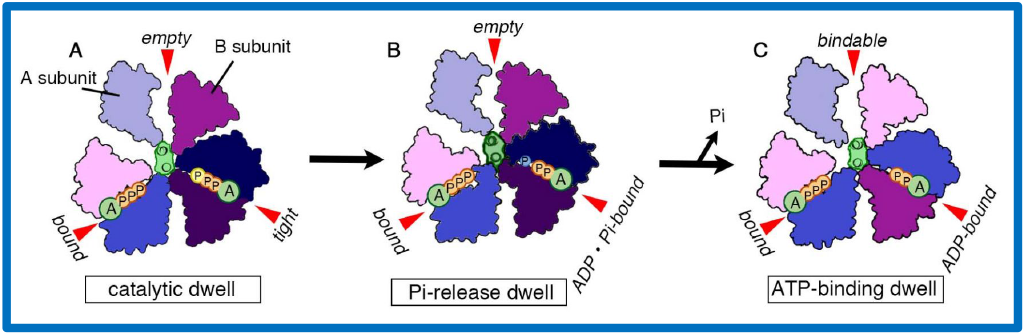
Proposed model of the rotation mechanism of *E. hirae* V_1_-ATPase. **(A–C)** The structure models are based on the crystal structures of catalytic dwell (2_(AMP · PNP)_V_1_ in panel A), P_i_-release dwell (2_(ADP · AlF4)_V_1_ in panel B) and ATP-binding dwell (2_ADP_V_1_ in panel C). The ATP indicated with a yellow terminal P_i_ is committed to hydrolysis.

### Transition from catalytic dwell to P_i_-release dwell is spontaneous after ATP hydrolysis

We replaced the AMP · PNP in the *tight* pocket of the catalytic dwell (3VR6) model with an ADP · P_i_ (P_i_ modeled as H_2_PO_4_^-^), and simulated the system in explicit solvent using allatom MD for 500 ns. Each simulation was replicated thrice, wherein the *empty* site was left nucleotide-free and the *bound* site included in ATP (list of simulations provided in Table S1). Similar simulations were also performed with the *tight* AMP · PNP replaced by an ATP^25^. To construct these simulation models, the ATP, ADP and AMP are aligned based on the geometry of the adenine ring and the first phosphate moiety^26^.

As illustrated in **Fig. 4**A, prior to hydrolysis, the ATP remains bound to the G235, G237, K238, R262 and F425 residues of the A subunit of the *tight* pocket and the Arg-finger R350 residue of the B subunit (denoted A^t^ and B^t^, and a similar site-wise nomenclature is followed for all other AB pairs). Consequently, the allosteric communication between the A^t^ and B^t^ subunits is relegated by the interface-bound ATP (**Fig. 4**C). On breaking the covalent bond to the terminal phosphate, the ADP stays connected to the A subunit, while the P_i_ interacts primarily with the Arg-finger of the B-subunit. Thus, the allosteric communication across the bound ATP is lost (**Fig. 4**D), resulting in a looser ADP · P_i_-bound AB interface. This looseness of the ADP · Pi bound interface is reflected in elevated fluctuations of the A^t^ subunit post hydrolysis (Fig. S2).

**Fig. 4:**
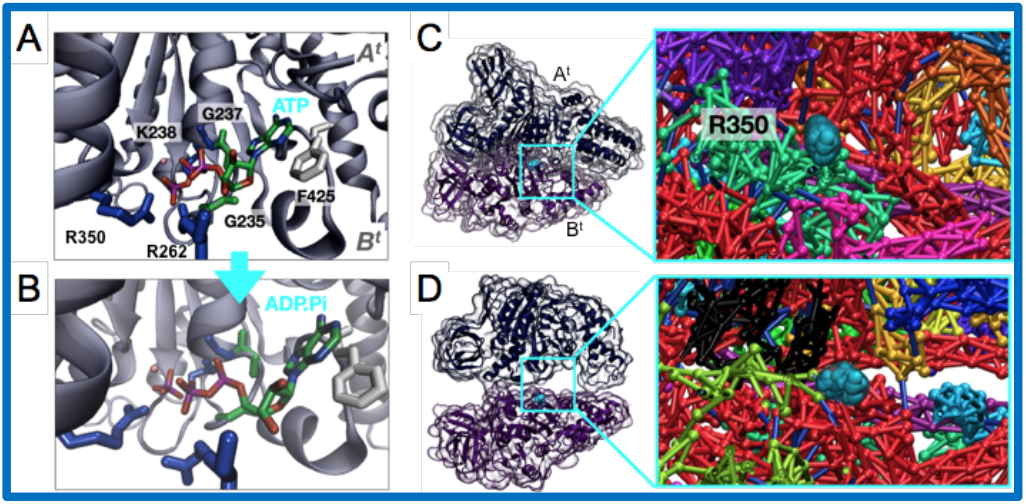
ATP hydrolysis breaks the dynamic coupling between A- and B-subunits. **(A)** ATP is bound to the binding pocket, formed by G235, G237, K238, R262, and F425 of the A-subunit and R350 of the B-subunit, prior to hydrolysis. In this state, the A^t^-and B^t^-subunits are tightly coupled due to the presence of the ATP. **(B)** Upon hydrolysis and prior to release of the inorganic phosphate (P_i_), the ADP and P_i_ moieties interact primarily with the A- and B-subunits, breaking the tight coupling seen in panel A. **(C)** Network model showing allosteric communication between the A- and B-subunits before hydrolysis, with R350 of the B-subunit highlighted in blue. Tight binding of the subunits gives rise to a strong network. **(D)** Network model showing allosteric communication between the A- and B-subunits after hydrolysis, with R350 of the B-subunit highlighted in blue. Breakage of the tight coupling between the subunits results in a weaker allosteric network.

The RMSD between the simulated ADP · P_i_ bound models and the P_i_-release dwell X-ray structure (i.e. the 2_(ADP · AlF4)_V_1_ model) is peaked ∼1.4 Å, which is lesser than the 1.7 Å RMSD between the catalytic dwell (i.e. the 2_AMP · PNP_V_1_ model) model (PDB: 3VR6) and our simulated ADP · P_i_ bound model. Though small, the differences between the ADP · P_i_ bound model and the catalytic dwell model are statistically significant (Fig. S3). This trend is amplified when we compare RMSD values between the simulated ADP · P_i_-bound model with that from the ATP-binding dwell following the release of P_i_ (PDB: 5KNB). Our simulated ADP · P_i_-bound model is found to be deviated from the 5KNB model by 2.3 Å, which is more than its deviation from the catalytic dwell seen in Fig. S3. Noting that RMSD between the *tight* forms of 3VR6 and 5KNB is ∼3 Å, we find that the V_1_ models with ADP · P_i_ bound to A^t^B^t^ is close to and yet distinct from the structure of the catalytic dwell, and is also deviated from the ATP-binding dwell, justifying thus its own dwell state.

MD simulations of the V_1_ motor after ATP hydrolysis therefore spontaneously deforms the structure from the catalytic dwell to the ADP · AlF_4_^-^ bound model, suggesting that indeed the crystal structure represents a state following ATP hydrolysis but preceding product release. The simulated ADP · P_i_-bound model also possesses remarkable similarity with structures from our previous string simulations^25^, wherein we found that the presence of ADP · P_i_ in the A^t^B^t^ pocket, entraps the V_1_ rotor in a deep energy minimum and inhibits further rotation of the central stalk (Fig. S4), prior to product release. Altogether, the similarity between the ADP · P_i_-bound structures derived from the current set of MD simulations, the ones from our previous string simulations showing rotational inhibition, and the ADP · AlF_4_^-^ bound X-ray model, as well as their common difference from the ATP-bound A^t^B^t^ (in catalytic dwell) and ADP-bound A^t^B^t^ (in ATP-binding dwell) suggests the identification of a new P_i_-release dwell.

Worth noting, the MD simulations began with a model of the catalytic dwell, namely 3VR6 (2.6 Å resolution), and arrived at models similar to the crystal structure of the P_i_-release dwell. This computational result suggests on one hand, a minimal impact of initial model bias on our inferences, while on the other, biological relevance of the 2_AMP · PNP_V_1_ model despite its 3.8 Å resolution.

### Transition from P_i_-release dwell to ATP-binding dwell involves two different pathways

If the hydrolysis product remains in the binding pocket, the rotation of the central stalk is hindered and the motor is inhibited^27^. Thus, the next step after hydrolysis is considered to be the release of the ADP and P_i_ products, which is suggested as the rate determining bottleneck of the rotatory cycle in V-type, as well as the more ubiquitous F-type ATPase motors^28^. Interaction energy analysis from the MD simulations revealed that the protonated inorganic phosphate has the weakest interaction with the A^t^B^t^ protein pocket, while ADP has much stronger electrostatic interactions with this pocket (Fig. S5). Building on this initial insight, we performed 512 ns of replica exchange with solute tempering or REST2 simulation using 16 replicas (see Methods). This enhanced sampling simulation brought to light some initial stages of the product release, wherein the P_i_ to ADP distance increased from 4 Å to 8 Å, however a complete detachment of P_i_ was not observed due to the finite timescale of the computation (Fig. S6).

To determine a mechanism of product release we therefore resorted to well-tempered Funnel metadynamics simulations^29^. A vector suggesting the direction of P_i_ release was already identified in the REST2 simulations (arrow shown in **Fig. 5**). Two 0.4 and 2 μs metadynamics simulations were performed, one biased towards the central stalk akin to the REST2 results, denoted *inward* direction, and another biased in the opposite *outward* direction. The outward pathway led to complete detachment of the P_i_ and its release in the solvent, while along the inward path the P_i_ did detach from the binding pocket, but found a transient secondary site close to the central stalk. After the P_i_-release from the outward pathway, the RMSD between our simulated ADP-bound A^t^ state (relaxed with an additional 100 ns of MD) with that of the ATP-binding dwell 5KNB is ∼1.5 Å (Fig. S3).

**Fig. 5:**
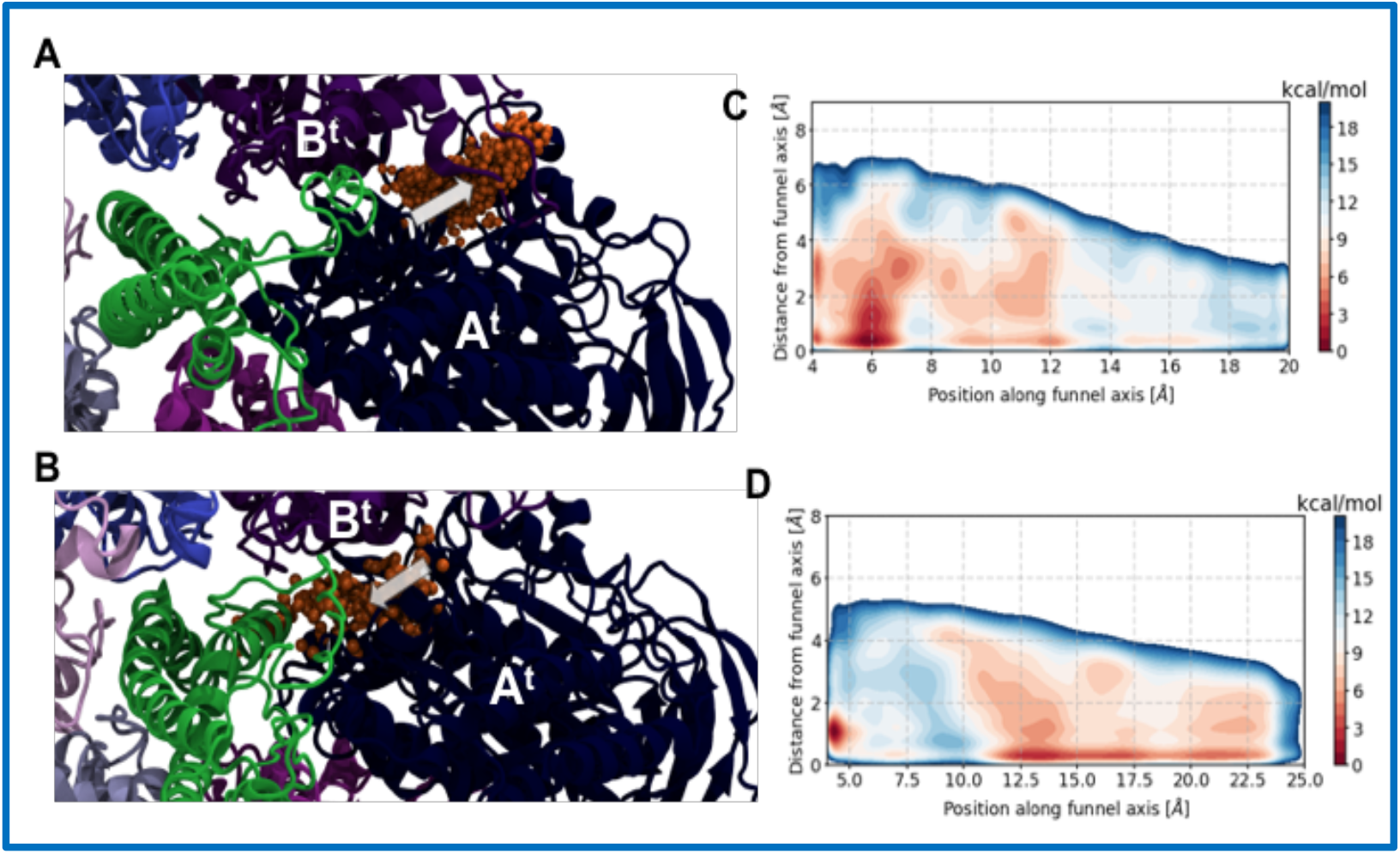
Pathways of P_i_ release. The two simulated pathways for phosphate release are either **(A)** *outwards* (away from the stalk) or **(B)** *inwards* (towards the stalk). The first event of phosphate release captured using funnel metadynamics is shown; P_i_ molecules are shown as orange spheres with the white arrow depicting the direction of egress. The free energy for the phosphate release computed by funnel metadynamics along the funnel and orthogonal axes is shown for **(C)** outward pathway and **(D)** inward pathway. In the vicinity of the binding pocket, the inward pathway is more constrained, and encompasses higher barriers following disengagement of P_i_ from the pocket. The outward P_i_-release pathway is energetically less expensive.

Free-energy profiles along both pathways have comparable energy barriers of height 6-7 kcal/mol for initial product release (Fig. S7). However, the inward release pathway has a number of unbinding intermediates (**Fig. 5** and Movie 1), which are missing from the outward release mechanism. The trapping of P_i_ in this intermediate suggests that in the solvated protein environment, overall, the inward release path out of the ATPase motor is slower than the outward release mechanism using Eq. 2 (see Methods). Evidently, the outward release takes 0.34 ms for the 18 Å pathway terminating the bulk solvent. In comparison, inward release requires 0.5 ms for the 15 Å pathway, which keeps P_i_ proximal to the stalk rather than the bulk solvent. However, the inward pathway requires less rearrangement of the product-containing AB interface, thus allowing for a small aperture for rapid P_i_ release from the pocket, after which, the P_i_ still remains non-specifically bound to parts of the stalk-A interface for a longer duration. The lifetime of the product release dwell is at least 0.5 ms. Therefore, we suggest that P_i_ is released before ADP, and the duration of the P_i_-release dwell will be controlled by the kinetics of the inward release pathway. Noting that single molecule experiments have now reached the time resolution of microseconds only very recently^8^, it is expected that this new dwell can also be seen in future experiments.

A transfer entropy analysis (see Methods) along the inward pathway further reveals that information is exchanged between the backbone RMSD of the A^t^B^t^ binding pocket and the reaction action coordinate vector of the product release (Fig. S8)^30^. We find that residues GLU261, ARG262 and PHE425 from subunit A and R350 from subunit B, that are implicated in ATP hydrolysis and/or binding^2^, are also involved in transferring information from the hydrolysis-site to induce larger scale conformational transition of the A^t^B^t^-pocket in the P_i_-release state into that of the ATP-binding state. In addition, distal hydrophobic residues VAL230, LEU291, ILE292 that are neither involved in the hydrolysis nor engage in ATP binding also participate in coupling the local event of product release with the global changes in conformation. Interestingly, coupling of the Arg finger R350 conformations with the global changes of the B subunit is 1.5-fold more pronounced in the ADP · P_i_-bound B^t^ site than in the B^e^ site. Based on this difference in sidechain information between the B^t^ and B^e^ sites, it is inferred that disengagement of the P_i_ from R350 will break the coupling between local and global conformations of the B subunit. Since the breaking of such allosteric information channels is energetically expensive^31^, the unbinding of P_i_ from the Arg finger is expected to offer a rate-determining barrier for the transformation of the P_i_-release dwell into the ATP-binding dwell.

According to Boyer’s ‘binding change’ model in the hydrolysis direction, any of the three catalytic sites on the enzyme first unbind ADP and/or phosphate in sequence and second, undergo a conformational change so as to intake and subsequently bind a new ATP^9^. Akin to the first step of this model, originally proposed for F-type ATPases, our results now show that even in the V_1_ motor, the enzyme unbinds phosphate and ADP in a sequence from the A^t^ sight, and not simultaneously. Next, to complete the second step of Boyer’s model for V-type ATPase i.e. to move from the P_i_-release dwell to the ATP-binding dwell of **Fig. 3**, it is necessary that information is transferred from the A^t^ site to the A^e^ site so a new ATP can enter the motor from the *e* site after the product is released from the *t* site. However, such information exchange between local ligand changes and global conformational changes (*ala* induced fit) is less feasible at the A^e^ site, where neither ATP or ADP · P_i_ binding is significant^2^. So, we probe protein-protein interface changes between the neighboring 120° apart *t* and *e* sites to elucidate how product release enables new reactant entry.

Post phosphate release, we find that the interaction between the A^t^ and the central stalk loosens, making the stalk more flexible. This additional mobility of the stalk also alleviates its interactions with the A^e^ and B^e^ (**Fig. 6**). Consequently, the solvent accessible surface area of the A^e^ pocket increases by ∼25%, allowing space for potential entry of a new ATP, and completing the P_i_-release dwell → ATP-binding dwell transition. The flexibility of the central stalk has been an issue of major contention in the F-ATPase area. While the crystallographers found no evidence of a deformed stalk in the molecular models, the single-molecule imaging and computational studies found evidence of a spatially dependent elastic modulus of the stalk^32^ in the F_1_ motor, indicating flexibility and potential deformation of the stalk during rotation. The central stalk of the V_1_-ATPase was already crystallographically shown to be deformed^5^. Here we find a design advantage of such stalk mobility that allows information transfer between P_i_ release from the *tight* pocket and reactant entry in the neighboring *empty* pocket.

**Fig. 6:**
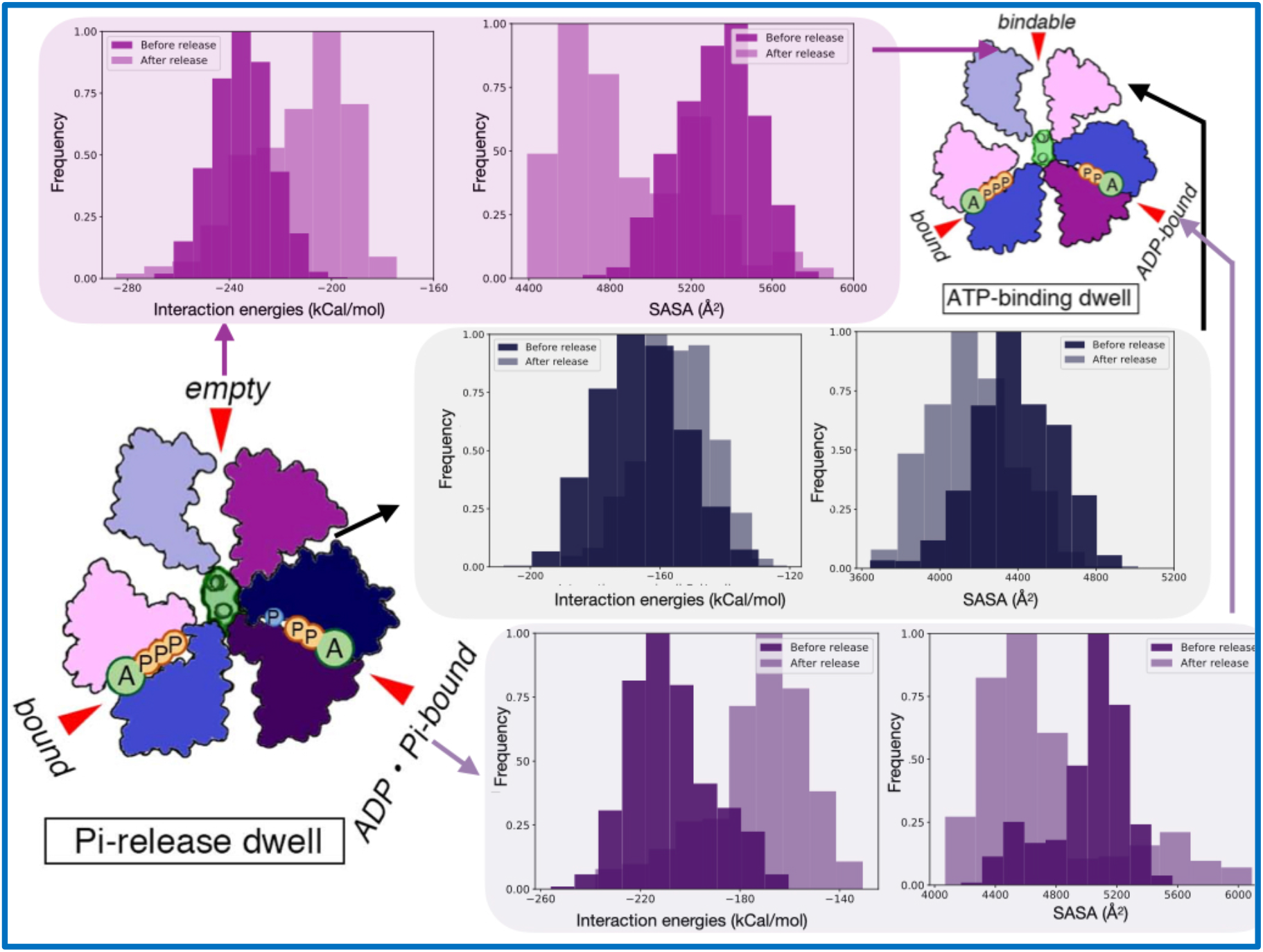
Interface energy analysis. Upon P_i_ release, interaction energy reduces (becomes less negative) between the stalk and subunit A^e^B^e^ (top panel, left), B^e^A^t^ (middle panel, left), and A^t^B^t^ (bottom panel, left). Concomitantly, the solvent accessible surface area (SASA) of the empty pocket (top panel, right), non-nucleotide binding pocket (middle panel, right) and the ADP · P_i_-bound pocket (bottom panel, right) undergoes an increase. This increase in SASA of the empty pocket allows it to accept a new ATP molecule to reset the rotatory catalysis cycle.

## DISCUSSION

Almost four decades after Boyer and Walker’s proposal of a rotatory catalysis mechanism in ATPases, we are still in the middle of molecular biophysics investigations resolving the details of the ATP activity^33^. Static snapshots determined using X-ray crystallography and MD simulation pictures of the V_1_ rotary motor from *E. hirae* are presented and compared in the schematic of **Fig. 7**. Simulation studies provide a complementary view of the rotation and ATP hydrolysis, by connecting the static intermediate structures during rotation. We had already established that after the *empty* form (in the catalytic dwell) changes to the *bindable* form (in the ATP-binding dwell), new ATP is bound to induce further conformational changes to drive the central stalk rotation, which appears to undergo a wringing movement during rotation to reset back to a new catalytic dwell^25^. The timescale of the rotation of the central stalk was determined to be of the order of 1.4 ms, but only after both the ADP and P_i_ were released from the *tight* pocket.

**Fig. 7:**
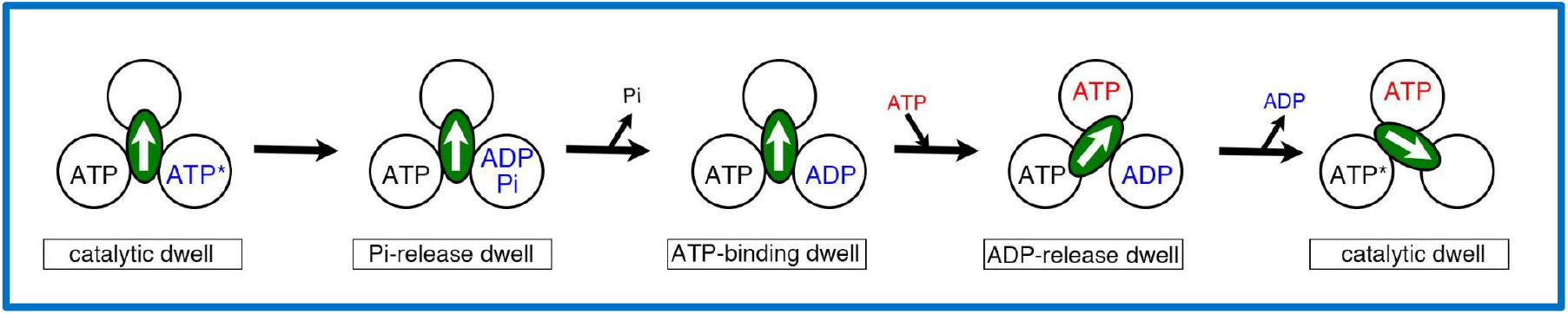
Proposed coupling scheme for ATP hydrolysis of *E. hirae* V_1_-ATPase. Each cycle in the figure represents the chemical state of the nucleotide-binding site from the cytoplasmic side. The central arrows in the ellipses represent the orientation of the central axis beginning from the twelve o’clock position, which corresponds to the catalytic dwell. ATP* represents an ATP molecule that is committed to hydrolysis. See Conclusion for additional details.

Here, we focus specifically on the V-ATPase’s product-release process. We find that the unique asymmetry of the A_3_B_3_ ring with three identical AB pairs facilitates P_i_ release and introduction of a new ATP into the hexameric motor prior to this stalk rotation. The ATP hydrolysis intermediate, that is trapped here using ADP · AlF_4_^-^, spontaneously relaxes an A^t^ pocket to marginally open conformation creating a P_i_-release dwell. This new intermediate in turn, facilitates the formation of a small aperture for releasing the P_i_ either internally towards or away from the stalk. Residues required for the ATP binding and hydrolysis are also the ones found responsible for coupling the P_i_-induced binding pocket reorganization with larger scale conformational transformations of the A-subunits. In particular, disengagement of P_i_ from the Arg350 finger residue is deemed a rate-determining step of the product release process. Noting that the Arginine finger is ubiquitous across many AAA+ ATP-hydrolyzing motors that use the so-called Walker motif^34^, the P_i_ -release mechanism presented here is generalizable to these systems.

The ADP-bound conformation formed after the P_i_ reduces the stalk-A subunit interaction, opening up a neighboring AB interface for accepting a new ATP. Through molecular simulations we find how the two ATP/ADP · P_i_-sites communicate with each other for synchronizing the protein conformations necessary for product release with the ones needed for ATP entry. Evidence of such multi-site conformational allostery was not included in the original binding-change model. Therefore, our joint structure-determination and computational work brings to light how the cooperativity between the AB proteins works inside a hexameric architecture to connect the P_i_-release dwell with the ATP-binding dwell.

Finally, comparing all the available kinetic data, we note that a 120° rotation back to the catalytic dwell takes ∼1.4 ms^25^, which is slower than the cumulative time of catalytic dwell → P_i_ release dwell → P_i_ release → ATP-binding dwell transition that takes no more than 0.5 ms. Given the turnover time of V_1_ in the hydrolysis direction is between ∼100 sec^-1^, the remaining molecular events associated with the ADP unbinding dwell can therefore be attributed to be the slowest along the entire rotary-catalysis cycle. Since ATP entry to a *bindable* pocket is spontaneous, through inspection of all the steps, we can now pinpoint that ADP-release is the rate-limiting step of V_1_-ATPase turnover. This finding is in line with single-molecule experiments that have implicated ADP accumulation and inhibition as the central cause for stoppage in the rotatory movement of V-type motors^8,35^. Now, our proposed mechanism of rotary-catalysis brings to light the molecular origins of the inhibition as well as activity of V_1_-ATPase.

## METHODS

### Protein Preparation

The A_3_B_3_ and DF complexes were expressed in *Escherichia coli* system, as described previously^36^. This system employs a mixture of plasmids containing the corresponding genes. A_3_B_3_DF complex (EhV_1_) was reconstituted and purified as follows^37^: purified A_3_B_3_ and DF complexes in buffer A (20 mM Tris-HCl, 150 mM NaCl and 2 mM dithiothreitol (DTT); pH 8.0) were mixed in a 1:4 molar ratio with the addition of MES (100 mM final concentration; pH 6.0) with 5 mM ADP and 5 mM MgSO_4_. After 1 h AlCl_3_ (1 mM final concentration) and NaF (5 mM final concentration) were added and incubated for 30 min at room temperature. V_1_-ATPase was purified using a Superdex 200 10/300 GL (GE Healthcare) column equilibrated with buffer B (20 mM MES, 10% glycerol, 100 mM NaCl, 5 mM MgSO_4_ and 2 mM DTT; pH 6.5). Purified complex was concentrated with an Amicon Ultra 30 K unit (Merck Millipore).

### Measurement of ATP activity

ATP hydrolysis activities of the EhV_1_ were measured using ATP regenerating system^2,3^. ATP hydrolysis rates were determined in terms of the rate of NADH oxidation, which was measured for 1 min as a decrease in absorbance of 340 nm at room temperature, and the measurement was repeated three times. Protein concentrations were determined using the Pierce BCA Protein Assay Kit (Thermo Fisher Scientific) with bovine serum albumin as the standard.

### Protein crystallization and Structure determination

Crystals of EhV_1_ were obtained by mixing 0.5 μl of purified protein solution (17 mg ml^-1^ protein in buffer B) with 0.5 μl reservoir solution (0.1 M Bis Tris propane, 0.2 M NaF, 20.8% PEG-3350 ; pH 6.5), using the sitting-drop vapor diffusion method at 296 K. Crystals were soaked in the solution (0.1 M Bis Tris propane, 21% PEG-3350, 5 mM ADP, 1 mM AlCl_3_, 5 mM NaF, 3 mM MgCl_2_, 0.28 M NaCl, and 20% glycerol ;pH 6.5) for 1 min, mounted on cryo-loops (Hampton Research, Aliso Viejo, CA, USA), flash-cooled, and stored in liquid nitrogen.

The X-ray diffraction data were collected from a single crystal at a cryogenic temperature (100 K) on BL-17A (λ = 0.9800 Å) at the Photon Factory (Tsukuba, Japan). The collected data were processed using XDS^38^ software. The structure was solved by molecular replacement with Phaser^39^ as a search model for nucleotide free V_1_-ATPase (PDB ID number: 3VR5). The atomic model was built using Coot^40^ and iteratively refined using Phenix^41^ and REFMAC5^42^. TLS (Translation/Libration/Screw) refinement was performed in late stages of refinement. The refined structures were validated with RAMPAGE^43^. All molecular graphics were prepared using PyMOL (The PyMOL Molecular Graphics System, Version 2.1.1, Schrodinger, LLC, New York, NY, USA).

### Molecular dynamics simulations

The MD study is based on a crystal structure of the V_1_-rotor from *Enterococcus hirae* (PDB ID: 3VR6)^2^. For simulation purposes, the artificial ATP mimetics, namely, AMP · PNP, which are employed as inhibitors for isolating crystals, are replaced by real ATP molecules in the bound states (*b*), and ADP and an inorganic phosphate or P_i_ (modeled as H_2_PO_4_ ^−^) in the tight state (*t*) (see **Fig. 2**). Structural analyses^2^ have demonstrated that non-hy-drolyzable AMP · PNP can successfully mimic the ATP and ADP + P_i_ binding states in F_1_-ATPase; different ATP analogues produced in fact, similar binding conformations^27^. The simulation was performed for the entire V_1_-rotor.

The V_1_-rotor is solvated in a water box of size 170 × 170 × 190 Å^3^ with 150 mM NaCl; the simulation system size is 0.49 M atoms. This structure was solvated and ionized in VMD, wherein 168273 water molecules were added. After a 4000-step energy minimization, the system is thermalized to 300 K in 50 ps at 1 atm, employing harmonic positional restraints with a 1 kcal/mol/Å2 spring constant on heavy atoms. Keeping the same spring constant, a 1 ns equilibration in the isobaric–isothermal ensemble (1 atm at 300 K) is carried out, followed by a 4 ns canonical-ensemble simulation, gradually decreasing the spring constant to zero during the latter stage. All MD simulations in our study are performed using NAMD 2.14^44^ with the CHARMM36 force field with correction for overcompensation for left-handed helix and CMAP corrections. The CHARMM-compatible P_i_ force field parameters were developed and used by us in previous studies^23^.

### REST2 simulations

Initial structure of ATP-synthase was obtained from the AlF_4_-bound crystal structure (resembling intermediate 1’ of the ADP.P_i_ bound state (Fig. S4), previously determined using MD simulations^25^). This structure was then minimized, heated to 300 K, and equilibrated for 5 ns. REST2 simulation was performed using 32 replicas scanning a temperature range of 300 K to 3000 K, where the ADP.P_i_ was defined as the solute. Each of the 32 REST2 replicas was simulated for 16 ns to attain a steady exchange-rate of 40-80% (Fig. S6).

### Funnel metadynamics and free energy computations

In order to accelerate the unbinding/binding of Pi,well-tem-pered Funnel-metadynamics (FM)^29^ was performed for 2000 ns and 400 ns for dissociation away or towards the stalk. The FM simulations were performed in an NVT ensemble using BioSimspace implementation in OpenMM. Following the OpenMM flavor of FM, a history-dependent bias was applied on two orthogonal CVs namely projection and extension that define the funnel potential. The projection CV which is collinear to the funnel axis is defined by a vector connecting the center of mass Ca of ILE322, GLU261, ASP 329 (Po) and GLU126. GLU275, ASP364 (Px) for dissociation away from the stalk and GLY353, PRO 268 for dissociation towards the stalk. The extension CV on the other hand, is orthogonal to the projection CV and is restrained by a sigmoidal restraint function defined as follows.

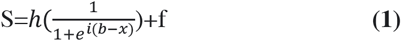

The parameters used to define the sigmoidal restraints are described in Table S2. For the FM, Gaussians hills with an initial height of 0.6 KJ · mol^-1^ was applied every ps, scaled by a WT scheme using a bias factor of 30 The hill widths chosen for the projection and extension CVs are 0.25 and 0.3 Å, respectively.

### Kinetics Calculations

Kinetics calculations provide an estimate of the mean first passage time, *τ*, which is the inverse of the rate constant *k*. Rate estimates and the mean free passage time was calculated from the free energy profile obtained from FM simulations using the following expression derived by Szabo et. al^45^.

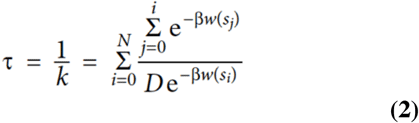

Here, the Pi dissociation pathway *x* is discretized into points *i, i+1*,….*k*,..*J. D*(x) represents the diffusion coefficient at point *x* and *s(x)* is the free energy at point *x* in the pathway. In our kinetics calculation *D(x)* was considered to be constant and was set to be 1.5 Å^2^/ns and 5.10 Å^2^/ns for dissociation towards and away from the stalk on the basis of previous diffusion calculation of Okazaki et.al^18^.

### Allosteric network analysis

The allosteric network analysis was performed with the NetworkView tool on VMD. The final 100 ns of the MD trajectories were strided by a factor of 10, creating nearly 1000 frames. These frames were analyzed by CARMA to create a covariance matrix, following which the allosteric networks are created at the C_α_-level of details, and the community substructures are analyzed using the Girvan-Newman method and default NetworkView parameters on VMD.

### Mutual information analysis

Given two random variables *X* and *Y*, mutual information is an information theory metric that quantifies the interdependence between *X* and *Y*. Mutual information (MI) is commonly expressed as

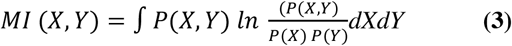

Here *P(X,Y)* is the joint probability distribution of X and Y and P(X) and P(Y) are their respective marginals. Inorder to understand the correlation between protein dynamics and ligand release MI was computed between the rmsd of protein backbone and position of Pi along the release pathway. In this case, *P(X,Y)* is the joint probability distribution of RMSD of the protein backbone and position of the ligand along the release pathway. We have in-house code to perform this analysis.

## DATA AVAILABILITY

The atomic coordinates and structure factors of 2_(ADP · AlF4)_V_1_ have been deposited in the Protein Data Bank under the accession code 7VW7.

## ASSOCIATED CONTENT

Supporting Information

The Supporting Information is available free of charge on the ACS Publications website:

- Additional analyses of the trajectories.
- Movies.

## ACKNOWLEDGMENTS

The synchrotron radiation experiments were performed at Photon Factory (proposals 2012G-132). We thank the beamline staff at BL1A and BL17A of Photon Factory (Tsukuba, Japan) for help during data collection. This research was supported in part by the Japan Agency for Medical Research and Development (AMED) under Grant no. 21fk0108092 (T.M.) and by a Grant-in-Aid for Scientific Research from Japan Society for the Promotion of Science (JSPS) under Grant no. 18H05425 (T.M.). Singharoy acknowledges the CAREER award from NSF (MCB-1942763). This work used the Extreme Science and Engineering Discovery Environment (XSEDE), which is supported by National Science Foundation grant number ACI-1548562.

## NOTES

The authors declare no competing financial interest.

## Supporting information

**Fig. S1.**
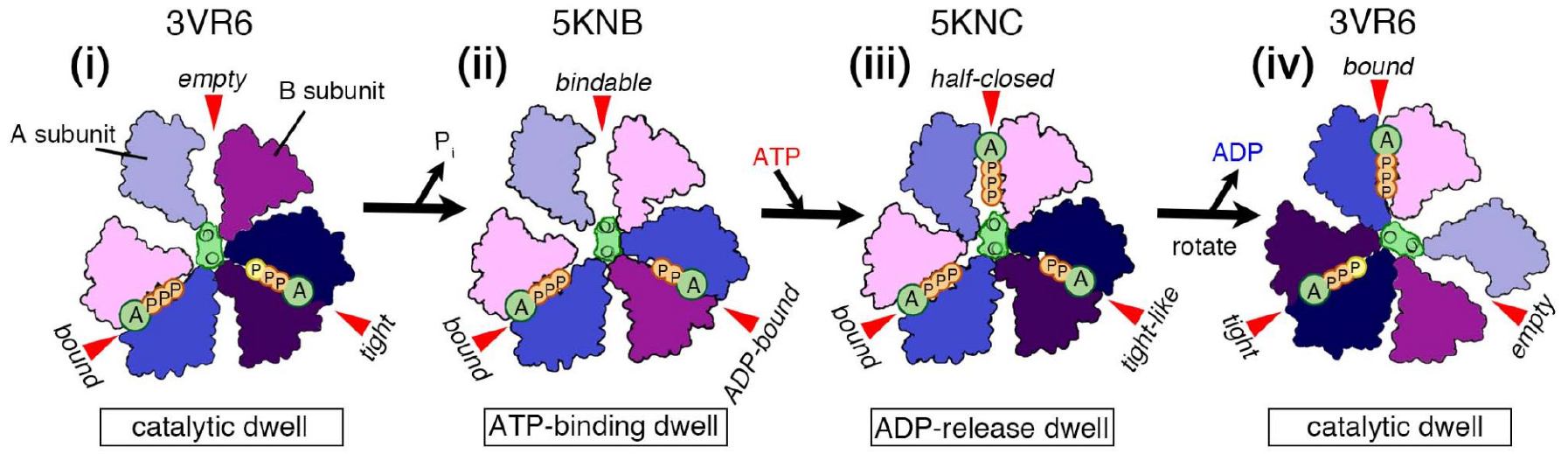
The model of the rotation mechanism of *E. hirae* V_1_-ATPase. The structure models are based on the crystal structures of catalytic dwell (bV_1_; **i**,**iv**), ATP-binding dwell (2_ADP_V_1_; **ii**), and ADP-release dwell (3_ADP_V_1_; **iii**), view from the cytoplasmic side. PDB ID numbers are shown above the structures. ATP indicated as a yellow ‘P’ in (**i**) and (**iv**) represents an ATP molecule that is committed to hydrolysis. The model starts from the catalytic dwell. ATP bound to the ‘*tight*’ form is hydrolyzed to ADP and P_i_, and the P_i_ release induces the conformational changes to the ATP-binding dwell. ATP binding at the ‘*bindable*’ form in the ATP-binding dwell state, induces the conformational changes to the ADP-release dwell. ADP release from the ‘*tight-like*’ induces the DF stalk rotation, and the consequent conformational changes to the catalytic dwell. See text for additional details.

**Fig. S2.**
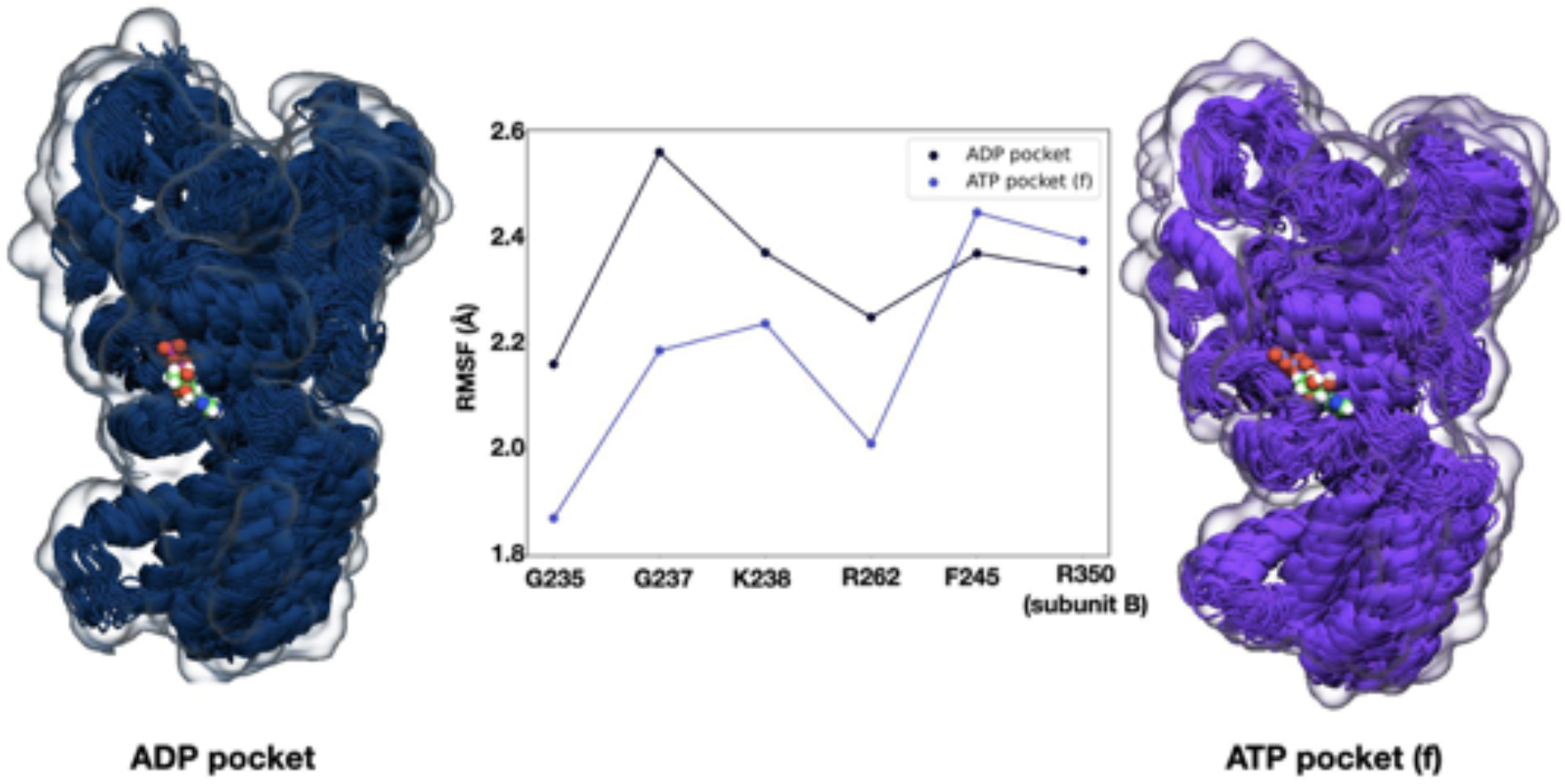
P_i_ release results in a looser AB interface. Root Mean Square Fluctuation (RMSF) of G235, G237, K38, R262, F245 of subunit A and R350 of subunit B, forming the ADP/ATP-binding site, shown for the ADP.P_i_-bound A^t^ pocket (dark blue) and ATP-bound A^t^ pocket (purple). The binding site is more rigid (lower RMSF) in the purple pocket. ADP/ATP-bound structures are shown in dark and light blue respectively.

**Fig. S3.**
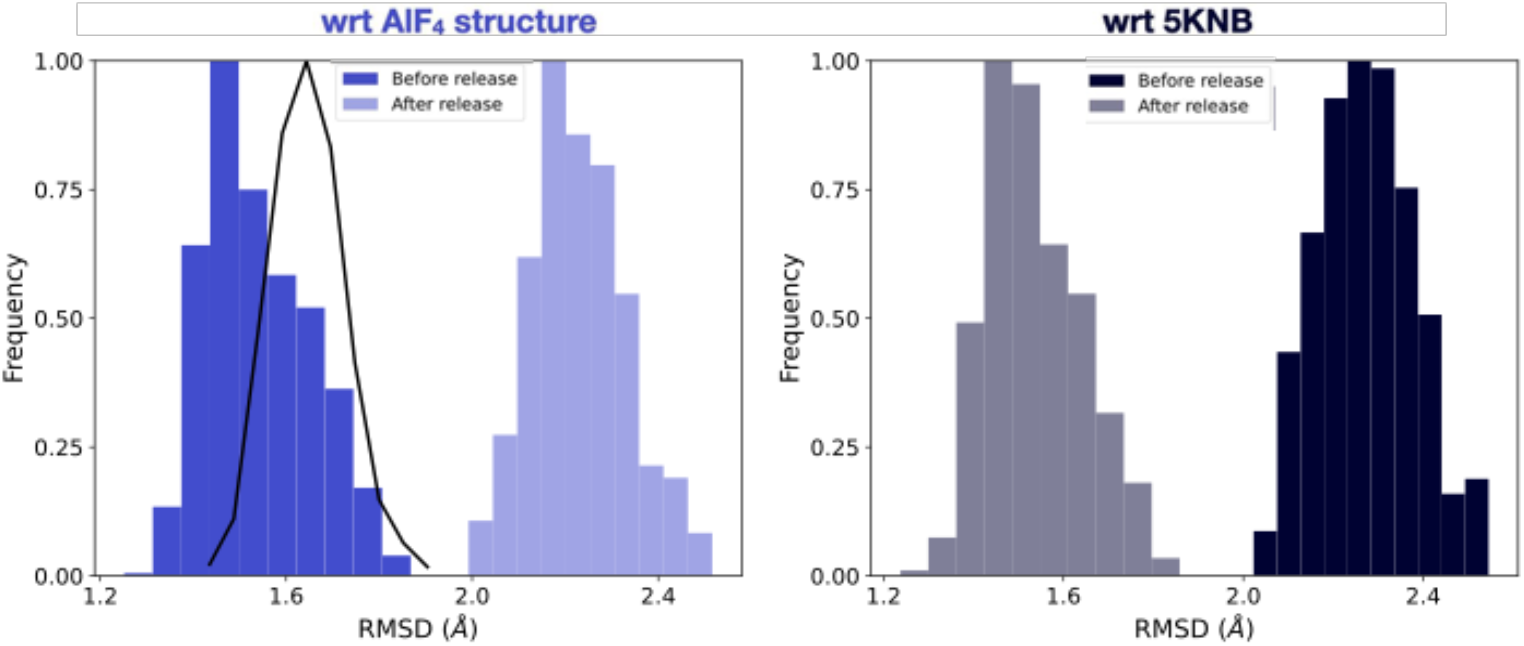
RMSD distribution of the simulated A^t^ pocket before and after P_i_ release measured with respect to the X-ray structure of the A^t^ pocket from newly discovered AlF_4_-bound “P_i_-release” dwell, and that with respect to the A^t^ pocket of previously resolved “ATP-binding” dwell (PDB: 5KNB in Fig. 1). Data for the pre-P_i_ released state is determined from MD and subsequent REST2 simulations; data for the post-P_i_ released state is determined by relaxing the final model from the 2000 ns-long outward pathway of the Metadynamics simulation further using 100 ns long regular MD. We observe that before the P_i_ release, the simulated pocket remains similar to the P_i_-release conformation, but after the P_i_ release, the same pocket drifts towards 5KNB. RMSD distribution of the ADP.P_i_-bound pocket with respect to the A^t^ pocket of the “catalytic dwell” of Fig. 1 is plotted a black solid line.

**Fig. S4.**
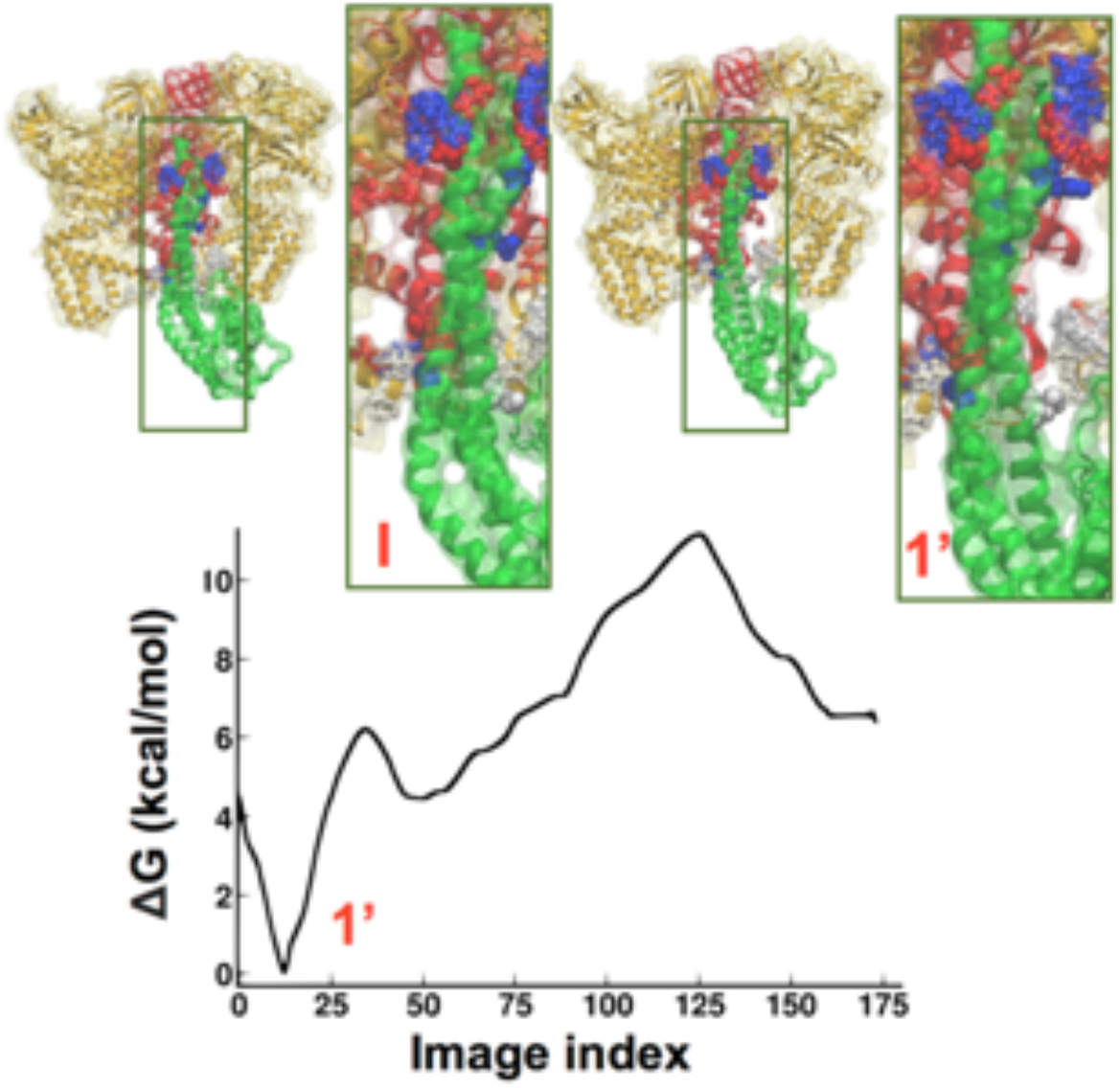
One-dimensional free energy profile characterizing the product (ADP + P_i_)–inhibited pathway of V_1_ – rotor structural transitions reveals that this process is endothermic [ref], and that the rotation is thermodynamically unfavorable in the presence of products bound to the empty site. An intermediate is observed along this pathway, denoted 1’, which resembles the P_i_-release dwell.

**Fig. S5.**
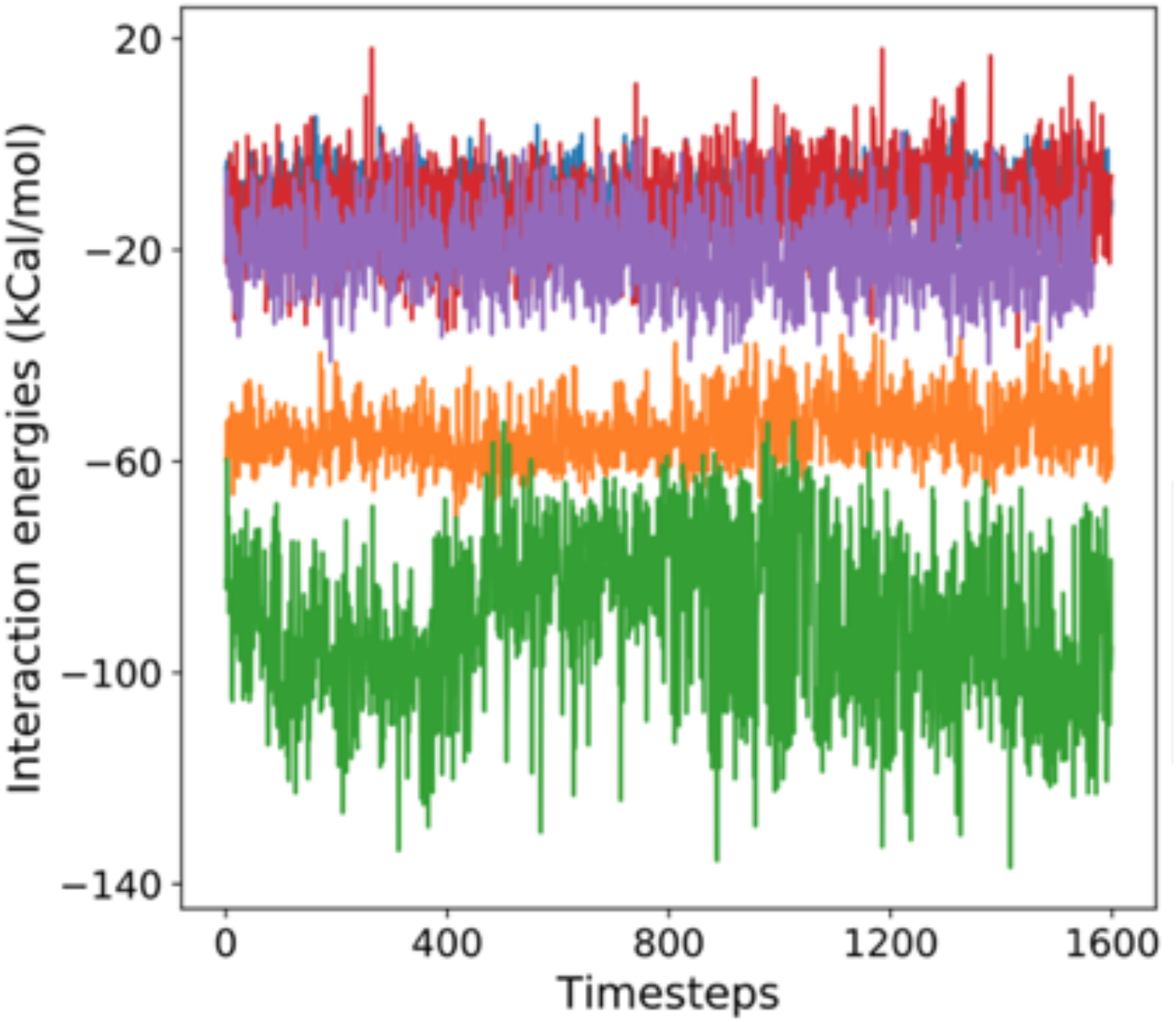
Interaction energies of P_i_ (blue) and ADP (orange) to A^t^, ATP in A^e^ pocket (red), and ATP in B^e^ pocket (purple) with the rest of the protein; timesteps are in ns cocatinatining three 500 ns equilibrium MD trajectory of the ADP.Pi-bound model of the P_i_-release dwell. Electrostatic interaction energy of ADP with the rest of the protein is also shown (green). ADP has more favorable (more negative) interactions compared to the other moieties, while the P_i_ has the weakest interaction amenable to its early release from the pocket.

**Fig. S6:**
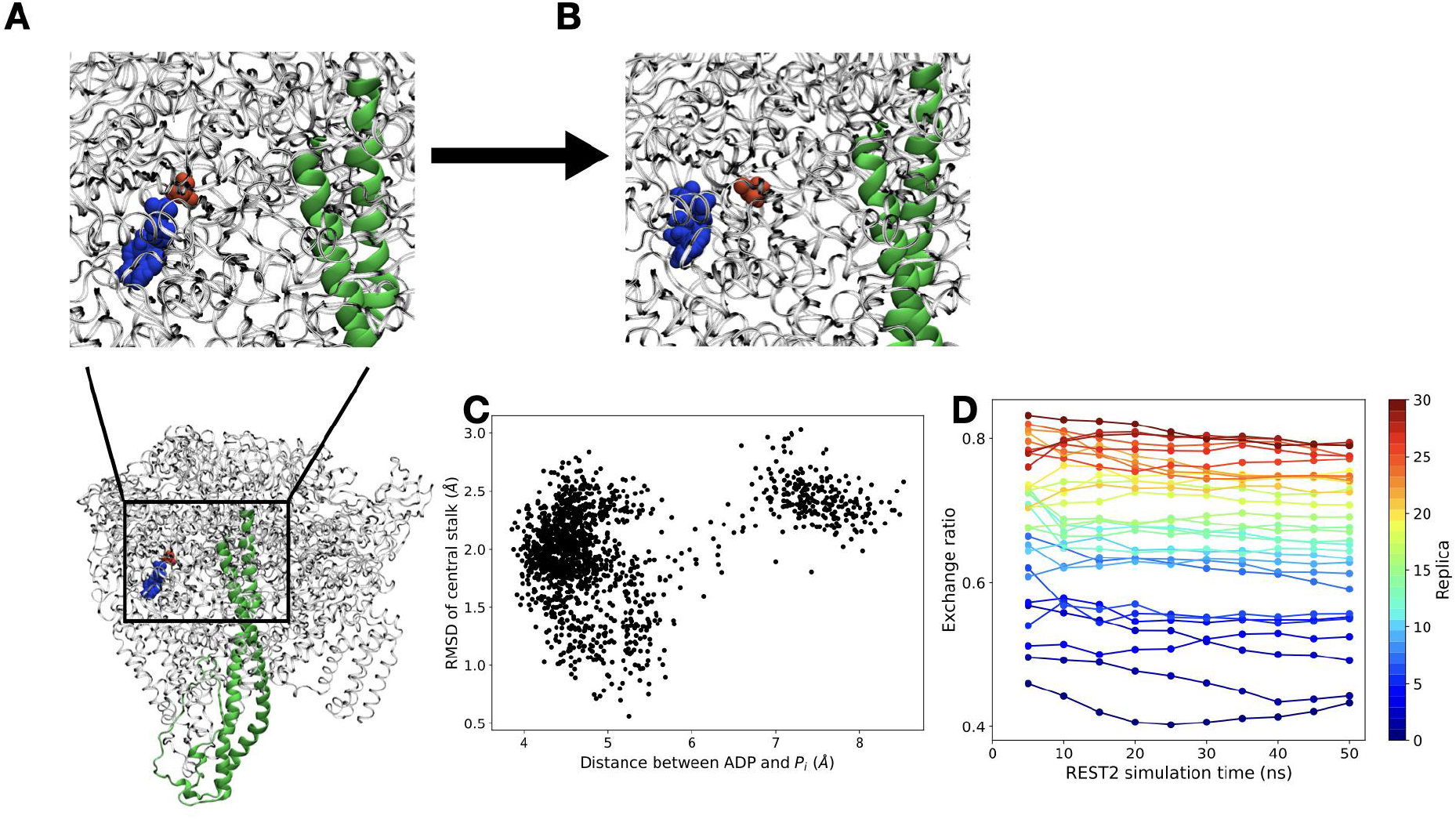
Rotation of the central stalk is coupled to the release of the P_i_ moiety. **(A)** ADP (blue) and P_i_ (red) before release. Zoomed-out view (bottom) shows the location of the ADP.P_i_ site wrt the central stalk. **(B)** ADP (blue) and P_i_ (red) after release. **(C)** RMSD of the central stalk w.r.t to initial conformation, plotted as a function of the distance between ADP and P_i_. **(D)** Convergence of REST2 simulation. Exchange rates between nearest neighbors of 32 replicas (31 rates in total) are shown over 50 ns of REST2 trajectory. Exchange rates vary between 40% and 80%, and stay essentially constant for a given replica.

**Fig. S7:**
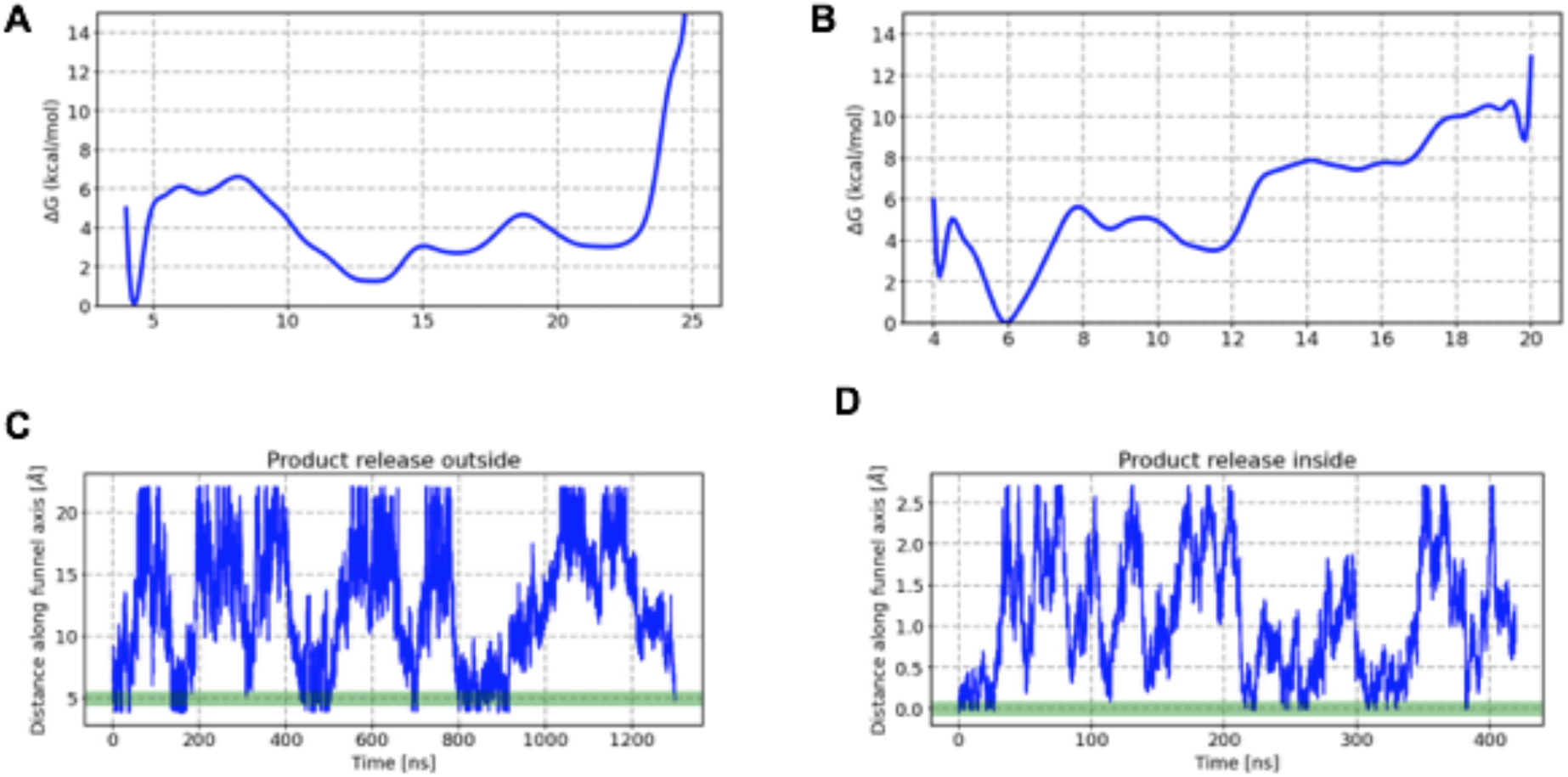
one-dimensional projection of the free energy of phosphate release. along the funnel axis for **(A)** *outward* pathway **(B)** *inward* pathway. Time series of the phosphate position along the funnel axis captured via funnel metadynamics for **(C)** outward release **(D)** inward release. Green band represents the crystal structure position of the phosphate.

**Fig. S8:**
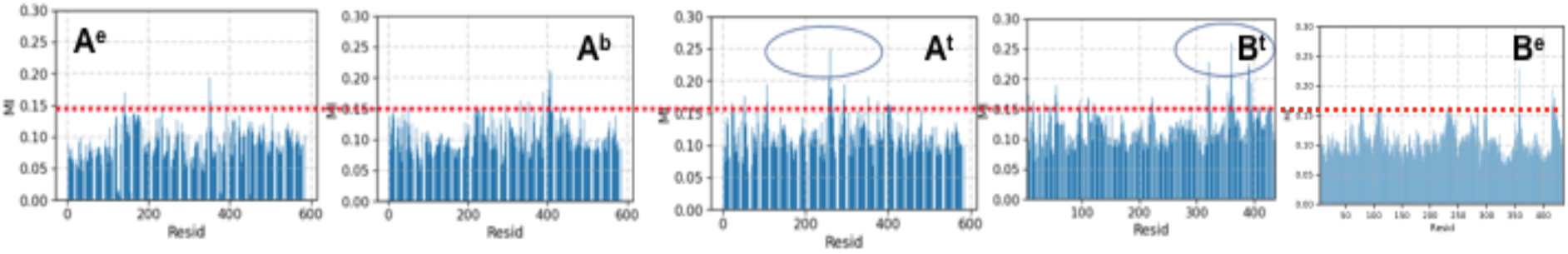
Bar Plot depicting the average mutual information. (MI) between phosphate position along the funnel axis and the backbone RMSD of residues in subunits A^e^, A^b^, A^t^, B^t^ and B^e^.

**Table S1:** List of simulations

| **Table S1:** List of simulations | System-size | Simulation length |
| --- | --- | --- |
| MD of catalytic dwell → P<sub>i</sub>-release dwell transition | 0.49 M | 3 replica x 500 ns/replica<br>(ADP.Pi bound to A<sup>t</sup>B<sup>t</sup>)<br>3 replica x 500 ns/replica<br>(ATP bound to A<sup>t</sup>B<sup>t</sup>) |
| REST2 of P<sub>i</sub>-release dwell | 0.49 M | 32 replica x 16 ns/replica |
| Metadynamics of P<sub>i</sub>-release | 0.49 M | 2000 ns (outward path)<br>400 ns (inward path) |

**Table S2:** Funnel Metady-namics Parameters

| **Table S2:** Funnel Metadynamics Parameters | Away from Stalk | Toward Stalk |
| --- | --- | --- |
| Beta-cent (b) | 15.0 Å | 15.0 Å |
| Wall width (h) | 6.0 Å | 6.0 Å |
| Wall buffer (f) | 1.0 Å | 1.0 Å |
| S-cent (i) | 12.0 Å | 15.0 Å |
| Lower Wall | 5.0 Å | 0.0 Å |
| Upper Wall | 20.0 Å | 26.0 Å |

**Movie S1:** Funnel metadynamics trajectory of the inward release pathway of P_i_ starting from the P_i_-release dwell approaching the ATP-binding dwell. Within our finite simulations, complete release of the P_i_ from the ATPase is not observed, though disengagement from the ATP-binding pocket is apparent.

**Movie S2:** Same as Movie S1, but for the outward release pathway, wherein the P_i_ after leaving the binding pocket completely disengages from the V_1_-ATPase.

## Notes

Supporting Information Placeholder

### Competing Interest Statement

The authors have declared no competing interest.

